# 9p21 Loss Defines the Evolutionary Patterns of Aggressive Renal Cell Carcinomas

**DOI:** 10.1101/2020.11.03.367433

**Authors:** Federica Carbone, Justin K. Huang, Luigi Perelli, Edoardo Del Poggetto, Tony Gutschner, Hideo Tomihara, Melinda Soeung, Truong Nguyen Anh Lam, Ruohan Xia, Cihui Zhu, Xingzhi Song, Jianhua Zhang, Kanishka Sircar, Gabriel G. Malouf, Alessandro Sgambato, Jose A. Karam, JianJun Gao, Eric Jonasch, Andrea Viale, Giulio F. Draetta, Andrew Futreal, Ziad Bakouny, Eliezer M. Van Allen, Toni Choueiri, Pavlos Msaouel, Kevin Litchfield, Samra Turajlic, Timothy P. Heffernan, Ying Bei Chen, Renzo G. DiNatale, A. Ari Hakimi, Christopher A. Bristow, Nizar M. Tannir, Alessandro Carugo, Giannicola Genovese

## Abstract

Dedifferentiation and acquisition of chromosomal instability in renal cell carcinoma portends dismal prognosis and aggressive clinical behavior. However, the absence of reliable experimental models dramatically impacts the understanding of mechanisms underlying malignant progression. Here we established an *in vivo* genetic platform to rapidly generate somatic mosaic genetically engineerd immune-competent mouse models of renal tumors, recapitulating the genomic and phenotypic features of these malignancies. Leveraging somatic chromosomal engineering, we demonstrated that ablation of the murine locus syntenic to human 9p21 drives the rapid expansion of aggressive mesenchymal clones with prominent metastatic behavior, characterized by early emergence of chromosomal instability, whole-genome duplication, and conserved patterns of aneuploidy. This model of punctuated equilibrium provides a remarkable example of cross-species convergent evolution.

**Significance:** To better understand the role of 9p21 in malignant progression, we generated a somatic mosaic GEMM of renal cancer, capturing the histological, genomic and evolutionary features of human disease. With this technology we demonstrated a critica role of 9p21 loss in metastatic evolution of RCC and provide a unique tool for testing new therapeutic treatments.

## Introduction

Aggressive vantians renal cell carcinomas (avRCC) are characterized by rapid clinical evolution and lack of response to therapy. The presence of metastatic disease, specifically, is associate with a particularly poor prognosis, however the mechanisms driving the emergence of malignant clones with metastatic competencies are still elusive (1). The lack of experimental models that faithfully recapitulate the clinical and pathological features of aggressive variants of RCC has significantly impacted our ability to understand the mechanisms underlying the malignant progression of this disease (2–5), to bridge these gaps in knowledge, we therefore developed an *in vivo* platform for the rapid generation of hight-throughput genetically engineered somatic mosaic mouse models (SM-GEMMs) of RCC.

## Results

Leveraging of CRISPR/Cas9-based genome editing method, we generated tissue specific somatic knock-outs of the tumor suppressor genes (TSGs) orthologs to commonly mutated genes in avRCC (Supplementary Fig. S1A-E) (6–8). Remarkably, such combinations consistently yielded indolent tumors characterized by low penetrance, long latancy, and limited invasive potential, with histopathological features of well differentiated papillary and tubulo-papillary carcinomas (Supplementary Fig. S1F-I). In order to find out whether additional genomic events are required to drive metastatic dissemination, leveraging *in vivo* somatic chromosome engineering, we generated a set of single-guide RNAs (sgRNAs) targeting DNA sequences flanking a 40-kb region on murine chromosome 4 syntenic to human 9p21. 3 (4q^9p21^). Strikingly, somatic engineering of the 4q^9p21^ locus loss resulted in the emergence of rapidly fatal tumors with a prominent tendency for widespread systemic dissemination and extensive sarcomatoid differentiation (sRCC) (*P* < 0.0001) (Fig. 1A-E, Supplementary Fig. S1J). These features are consistent with avRCC, mirroring their patterns of distribution and burden of metastatic spread (Fig. 1F-G) (9). Remarkably, analysis of two publically available datasets of RCC (TCGA, TRACERx) and an unpublished dataset of avRCC with clinical, pathological, and genomic annotations (MSKCC 2020), confirmed that the loss of genetic material on human chromosome 9p is associated with a significant increase in metastastic dissemination, presence of sarcomatoid features, and poor survival, particularly in *NF2*-mutant cancers (*P* = 0.0002 and *P* = 0.0048 for sarcomatoid features and metastatic disease, respectively) (Fig. 1H-K, Supplementary Fig. S2A-H, Supplementary Table S1) (10,11).

**Figure 1.**
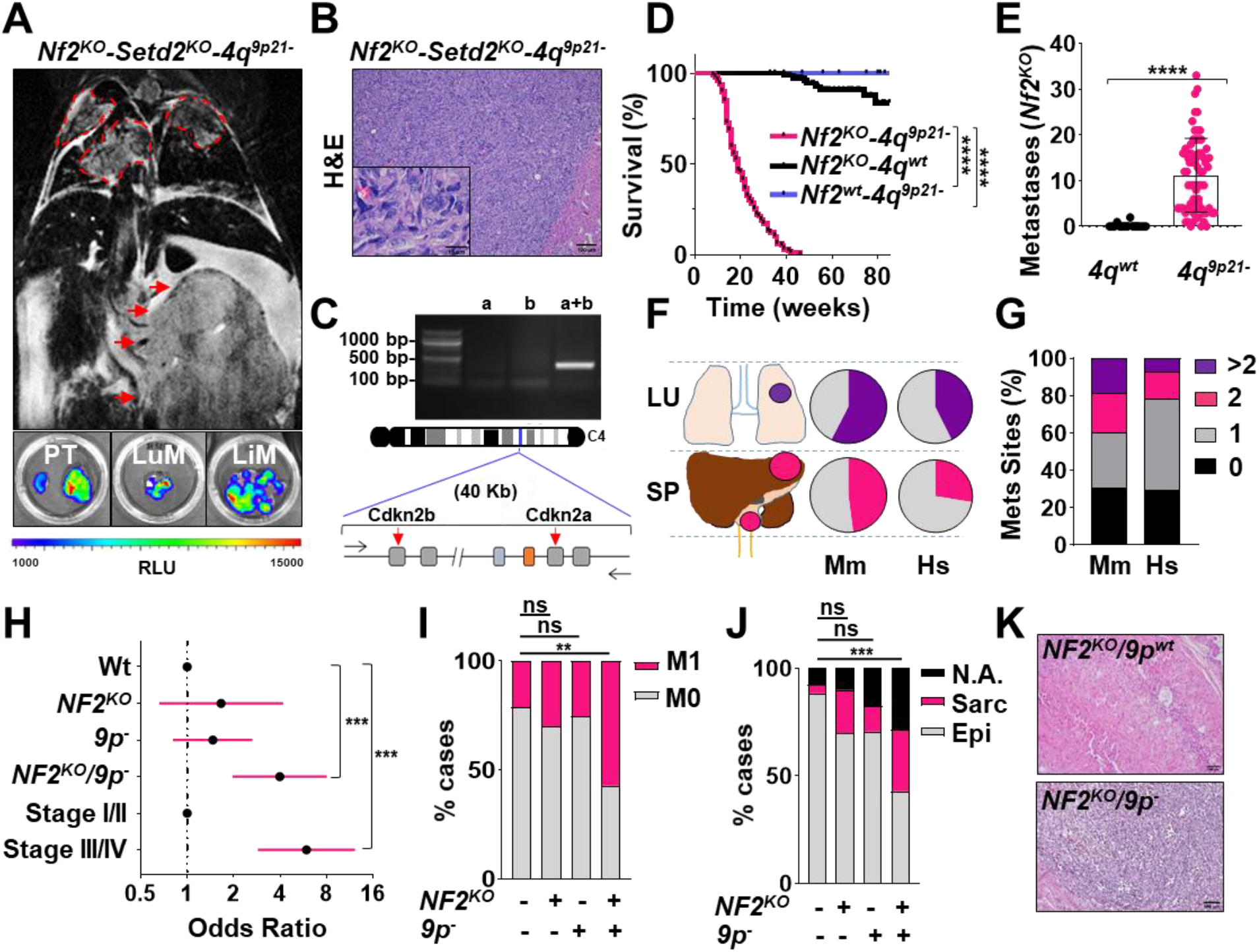
Somatic ablation of 4q^9p21^ drives metastatic progression in Nf2-driven SM-GEMM of RCC. **A)** *Upper panel*: representative coronal T2 MRI scan at 3 months post-transduction in *Nf2*^*KO*^-*Setd2*^*KO*^-*4q*^*9p21*-^ mice. Red arrows: primary tumor mass, red dashed lines: lung metastasis. *Bottom panels*: representative post mortem luminescence scans of mouse organs. PT: primary tumor, LuM: lung metastasis, LiM: liver metastasis. **B)** Representative H&E section of *Nf2*^*KO*^-*Setd2*^*KO*^-*4q*^*9p21*-^ tumors. Sarcomatoid features can be readily appreciated. **C)** *Upper panel*: deletion-specific PCR showing efficient deletion of the *Cdkn2a-b* locus on mouse 4q (single sgRNAs against *Cdkn2a* and *Cdkn2b* were used as negative controls). *Bottom panel*: schematic of the targeted murine 4q region. **D)** Kaplan–Meier survival analysis of *Nf2*^*KO*^-driven murine tumors with (N = 99) and without (N = 84) 4q^9p21^-targeting sgRNAs. Mice engineered with 4q^9p21^ alone do not develop tumors. **E)** Bar plot showing the metastatic burden in 4q^9p21-^(N = 69) vs 4q^wt^ (N = 15) models. **F-G)** Cross-species comparison of site-specific metastasis (F), disease burden and distribution (G). sRCC patients were used as reference. *Mus musculus* (Mm), N = 79; *Homo sapiens* (Hs), N = 199. **H)** Odds plot showing the enrichment of *NF2*^*KO*^/9p^−^ cases and stage III/IV features among MSKCC cohort patients. **I)** Bar chart showing the prevalence of metastasis features in *NF2*^*wt*^/*9p*^*wt*^, *NF2*^*KO*^/*9p*^*wt*^, *NF2*^*wt*^/*9p*^−^, and *NF2*^*KO*^/*9p*^−^ cases in the MSKCC cohort (N= 52, 10, 51, and 21, respectively) **J)** Bar chart showing the prevalence of sarcomatoid features in *NF2*^*wt*^/*9p*^*wt*^, *NF2*^*KO*^/*9p*^*wt*^, *NF2*^*wt*^/*9p*^−^ and *NF2*^*KO*^/*9p*^−^ cases in the MSKCC cohort (N = 52, 10, 51, and 21, respectively). **K)** Representative H&E stained images from two MSKCC cohort cases. *Upper panel*: *NF2*^*KO*^/*9p*^*wt*^; *bottom panel*: *NF2*^*KO*^/*9p*^−^. In the latter, sarcomatoid features are readily observed. N.S.: not significant, ** P < 0.01, *** P < 0.001, **** P < 0.0001 by log-rank (Mantel–Cox) test (D), two-tailed unpaired *t* test (E) and Fisher’s exact test (H, I, J). Error bars represent the standard deviation of biological replicates. Scale bar: 100 μm.

To characterize the genomic events driving rapid cancer evolution upon targeting the 4q^9p21^ locus, we performed multiregional whole-exome sequencing (WES) on tumor bearing mice and multiregional whole-genome sequencing (WGS) on selected cases, with an average coverage of 147X and 55X, respectively. The extent and distribution of somatic indels generated by the Cas9 endonuclease demonstrated patterns consistent with previous reports, showing high efficiency of *in vivo* editing, regardless of number and position of the sgRNAs engineered within a package of guides (Supplementary Fig. S3A-C) (12,13). This suggests high selective pressure for multiple clonal drivers, as previously described (10,12,14). Spatial variant allele frequency (VAF) analysis of specific engineered indels across disease sites further demonstrated the presence of one or a few dominant clones highly abundant within the primary tumors and enriched at secondary sites (Supplementary Fig. S3D-E). These data, along with conserved histopathological features across anatomical sites (Supplementary Fig. S3F), favor a monophyletic or oligophyletic origin hypothesis for this cancer model, with most of the disease burden established by the rapid fixation and expansion of clones with high fitness (15). In contrast with a relatively low mutation burden [0.34 somatic, exonic mutations (VAF ≥ 0.1 per Mb)], further characterization of acquired somatic variants revealed rampant chromosomal instability (CIN) with conserved patterns of chromosomal aberrations among differerent mice and lesions (Fig. 2A, Supplementary Fig. S4A-C). Remarkably, mouse/human synteny data evidenced cross-species convergent genomic evolutionary trajectories with recurrent losses affecting the mouse genome regions 12q and 16q, syntenic to human chromosomes 14q and 3p, respectively, and gains of the murine 11q and 5q regions, syntenic to human chromosomes 7 and 17 (Fig. 2B) (11). WGS analysis of selected cases coupled with karyotyping further revealed genome polyploidization, suggesting that clones undergoing one or multiple whole-genome duplication (WGD) events are selected during malignant progression, potentially creating a permissive background to tolerate high degrees of CIN (Fig. 2C-E, Supplementary Fig. S5A-D). Supporting this hypothesis, distribution analysis of heterozygous SNPs across the genome revealed that WGD events preceded the emergence of specific aneuploid events (Supplementary Fig. S5E). Further interrogation of TCGA, TRACERx and MSKCC RCC datasets confirmed a significant association between 9p loss, WGD, and high aneuploidy score (Supplementary Fig. S6A-C, Supplementary Table S1) with conserved patterns of genome evolution as demonstrated by the significant co-occurrence of 9p/14q losses (Fig. 2F, Supplementary Fig. S6D) (11,16). These results suggest that the loss of 9p is conductive to aneuploidy, promoting the rapid emergence of metastatic-competent clones (17).

**Figure 2.**
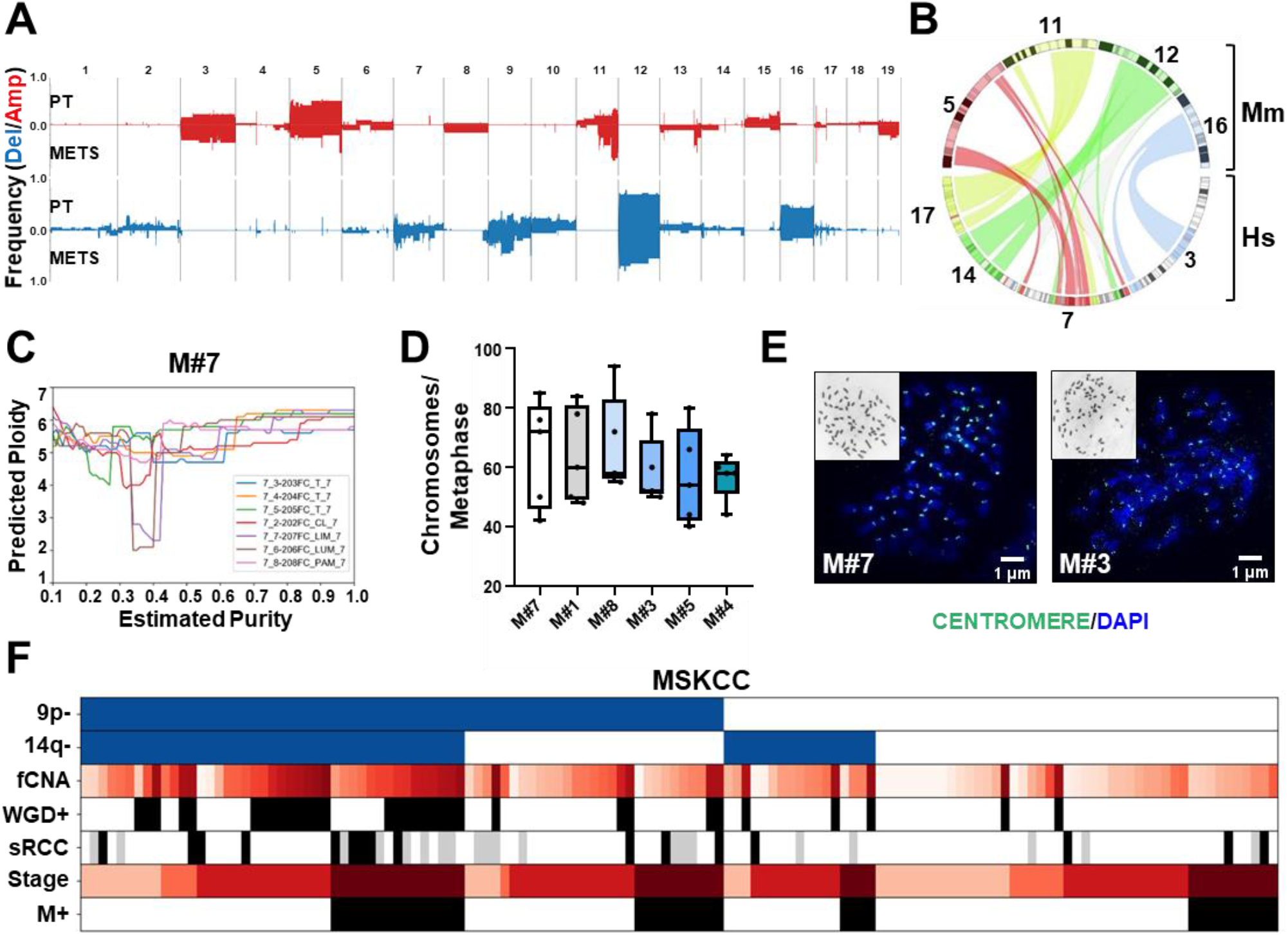
Convergent pattern of aneuploidy in murine and human aggressive RCC. **A)** Summary of segment-level amplification or deletion frequency across murine primary tumors or metastatic lesions as determined by GISTIC2. **B)** Circos plot of the human to mouse synteny map for chromosome regions significantly altered in SM-GEMM tumor-bearing mice generated by the SynCircos function of Synteny Portal (59,61). Remarkably similar patterns of syntenic chromosome gains and losses can be readily appreciated within TCGA dataset (51). **C)** Most probable ploidy by log posterior probability at given sample’s cellularity as predicted by Sequenza from WGS data (respesentative mouse #7). **D)** Chromosome counts in sRCC SM-GEMM–derived short-term cultures. Malignant cells are characterized by prominent polyploidy (N = 3/line tested). **E)** Representative co-staining of chromosomes (DAPI) and centromeres (ACA) in representative nuclei in metaphase from short-term cultures established from *Nf2*^*KO*^-*Setd2*^*KO*^-*Trp53*^*KO*^-*4q*^*9p21*^ tumor-bearing mice. Insert: G-banding analysis. **F)** Clinical and genomic annotation of specific features across the MSKCC RCC cohort (N = 134). Pairwise patients characteristics statistics are in Supplementary table S1. Error bars represent the standard deviation of technical replicates (D). Scale bar: 100 μm.

Accordingly, multiregional sequencing (MRS) analysis of primary tumors and metastases displayed the rapid truncal emergence of recurrent copy number variation (CNV) events (12q-, 16q-, 5q+, 11qE+) and a dramatic bias toward the accumulation of private over truncal single nucleotide variants (SNVs) (Fig. 3A-C, Supplementary Fig. S7A). This observation indicates that CIN emerges relatively early during the natural history of the disease in highly conserved patterns preceding exponential clonal expansion and metastatic seeding (Fig. 3D-E).

**Figure 3.**
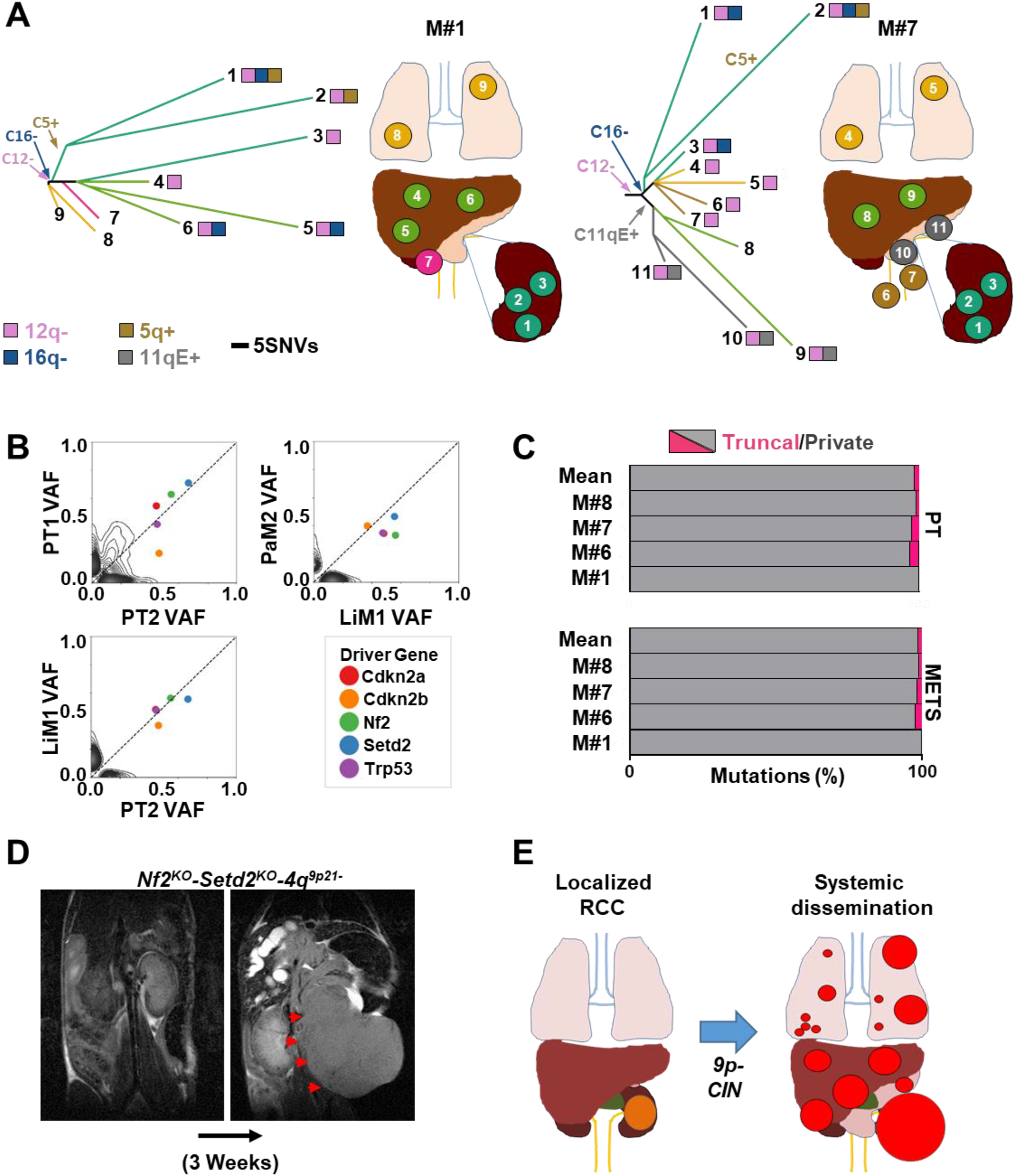
Aneuploidy and early clonal dissemination are hallmarks of aggressive RCC evolution. **A)** Representative phylogenetic reconstruction from MRS analysis of two advanced cases with metastatic disease, using sample progression trees based on presence or absence of somatic events. **B)** Density plots displaying the VAF of observed somatic mutations detected by WGS between selected representative samples (mouse M#7). Engineered events are reported in color. **C)** Bar charts showing the percentage of private and truncal somatic events at primary (*upper panel*) and metastatic sites (*bottom panel*) in tumor-bearing mice with advanced disease as determined by MRS. **D)** Coronal T2 MRI scan of Nf2KO-Setd2KO-4q9p21-tumor showing explosive patterns of disease progression. Red arrows: primary tumor mass. **E)** Schematic summary of sRCC evolution proposing a central role of 9p loss in driving rapid disease evolution and systemic spread.

## Discussion

Altogether, these results establish functional proof of the central role of 9p loss in determining aggressive patterns of disease evolution in RCC. Integrated genomic analysis of human renal cancers has highlighted a robust association between 9p loss, the presence of complex karyotypes and aggressive clinical behavior, across pathological and molecular subtypes of RCC, but there are still limited functional data supporting a role for 9p loss in malignant progression (10,11,18–20).

Despite of the crucial role of proper research platforms in understanding cancer pathophysiology and performing preclinical test of new therapeutic options, there is only an handful of renal cancer models, none of which showing consistent metastatic potential (21,22). In the present study, by engineering 9p21 loss *in vivo*, we generated the first somatic mosaic model of aggressive and metastatic RCC and captures the evolutionary features of human tumors. This platform ensures a rapid, high-throughput generation of different subtypes of RCC, harboring an intact immune system and therefore enabling future studies looking at the characterization tumor immune response, changes in tumor microenvironment and effects of treatment with immunotherapy agents.Here, we demonstrated that the loss of the murine ortholog of 9p21 triggers the explosive expansion of aggressive sub-populations, with mesenchymal features and prominent metastatic behavior. This results, strongly suggest that 9p21 restrains the emergence of malignant clones with invasive potential in renal cancer. Furthermore, whole exome and genome sequencing analysis provides insight into the patterns and tempo of genetic evolution in 9p-driven tumors, revealing early emergence and rapid selection of clones defined by whole-genome duplication events, chromosomal instability, and conserved patterns of aneuploidy. These features are in line with a model of punctuated equilibrium, where bursts of macroevolutionary events driving explosive patterns of tumor growth are followed by periods of neutral evolution and stasis *(23)*. Remarkably, mouse/human synteny data evidence cross-species convergent evolutionary trajectories, characterized by the spontaneous emergence of recurrent chromosome losses and gains. These findings point towards conserved evolutionary bottlenecks, shaping the natural history of RCC in both human and experimental models.

In conclusion, our somatic CRISPR-based chromosome engineering studies establish an efficient model recapitulating the main features of human cancer and functionally demonstrate a critical role for 9p21 loss in the metastatic progression of aggressive RCC. Taken together, these findings may lead to better understanding of cancer evolution and pave the road for new therapeutic strategies, providing substantial contribution to the advancement of the field.

## Methods

### Mice

The Pax8^Cre^ strain was generated by Dr. Meinrad Busslinger and obtained through the Jackson Laboratory, Stock No: 028196 (24). The H11^LSL-Cas9^ strain was generated by Dr. Monte M. Winslow and obtained through the Jackson Laboratory, Stock No: 027632 (25). The Rosa26^LSL-TdT^ was generated in Hongkui Zeng’s laboratory and obtained through the Jackson Laboratory, Stock No: 007908 (26). The Rosa26^FSF-LSL-TdT^ was generated in Hongkui Zeng’s lab and obtained through the Jackson Laboratory, Stock No: 021875 (27). Rosa26^LSL-Luc^ mice were generated by William G. Kaelin and obtained through the Jackson Laboratory, Stock No: 034320 (28). Strains were kept in a mixed C57BL/6 and 129Sv/Jae background. Embryo collection was performed at E14. All animal studies and procedures were approved by the UTMDACC Institutional Animal Care and Use Committee. All experiments conformed to the relevant regulatory standards and were overseen by the institutional review board. No sex bias was introduced during the generation of experimental cohorts. Littermates of the same sex were assigned randomly to experimental arms.

### Single-guide RNA design and validation

Single-guide RNAs (sgRNAs) were designed with “GenScript CRISPR sgRNA Design Tool” (https://www.genscript.com/gRNA-design-tool.html?a=post). First, 5′-phosphorilated oligos were annealed and diluted 1:20. Then 1 uL of each annealed and diluted sgRNA was cloned in digested lentiCRISPR V2 (addgene #52961) according to Dr. Feng Zhang’s protocol https://media.addgene.org/cms/files/Zhang_lab_LentiCRISPR_library_protocol.pdf). NEB^®^ Stable Competent *E. coli* (C3040I) colonies resistant to ampicillin antibiotic selection were amplified, and presence of sgRNA was confirmed by Sanger sequencing. Positive clones were transfected individually in 293 cells along with vectors for lentiviral packaging production, PAX2 (addgene #12260) and PMD2G (addgene #12259). MCT cells were infected by the lentivirus carrying a specific sgRNA and selected for puromycin resistance. Cut efficiency of sgRNA was tested by T7 Endonuclease I (NEB #M0302L) assay on the DNA of infected cells, according to the protocol (https://www.neb.com/protocols/2014/08/11/determining-genome-targeting-efficiency-using-t7-endonuclease-i)..

#### Single-guide RNA sequences

Nf2: GTATACAATCAAGGACACGG, Setd2: CTCGGGTGAAAGAATATGCA, Trp53: GACACTCGGAGGGCTTCACT, Cdkn2a: GTGCGATATTTGCGTTCCGC, Cdkn2b: GGCGCCTCCCGAAGCGGTTC, Bap1: GAATCGGTCTTGCTACTGCA.

#### Primers list

Nf2 For: CCTGCTTGTCTGGGAAGTCTGT, Nf2 Rev: GTCTCACCAACTAGCCATCTTCC; Setd2 For: TTGATTGCTGAAGGGTGTAACTCA, Setd2 Rev: CTGGCCTCAAACTTCCTAAACAGA; Trp53 For: CCGCCATACCTGTATCCTCC, Trp53 Rev: GCACATAACAGACTTGGCTG; Cdkn2a For: AAGGGCAGGGTGTAGAGTAAC; Cdkn2a Rev: CAGGTGATGATGATGGGCAA; Cdkn2b For: GGAATTAAGTGCTGGGTTGGAG, Cdkn2b Rev: CAGGACGCTCACCGAAGCTA; Bap1 For: GCCAGAACCACGTCACCTTC, Bap1 Rev: CAGGCCACAGGCAACCTAAA.

### Recombinant DNA

Packages of two or more guide RNAs were designed according to the following scheme: EcorI restriction site – U6 promoter – gRNA1 sequence – gRNA scaffold – polyA – U6 promoter – gRNAn sequence – gRNA scaffold – polyA – AscI restriction site. The synthetic sequence was synthesized and assembled into the pEMS2158-FLEx-Flpo AAV vector (Genescript) into the EcorI and AscI restriction sites. The pEMS2158-FLEx-Flpo was generated by PCR amplification of FLEx(loxP)-FlpO from the pTCAV-FLEx(loxP)-FlpO vector (addgene # 67829) (29) and cloning into the AscI and BsrGI sites of the pEMS2158 vector (addgene #70119) (30). AAV PHP.eB (addgene #28304-PHPeB) carrying FLEX-GFP sequence was used for injections in Pax8^Cre/+^-Rosa26^LSL-TdT/+^ mice (31).

### Virus production

Plasmid DNA preparations were generated using endotoxin-free MIDI kits (Qiagen). Large-scale AAV particle production was outsourced to Vigene Biosciences (10^^13^ IU/mL). Viral preparations were stored in aliquots at −80°C.

### Imaging

A 7T Bruker Biospec (BrukerBioSpin), equipped with 35mm inner diameter volume coil and 12 cm inner-diameter gradients, was used for MRI imaging. A fast acquisition with relaxation enhancement (RARE) sequence with 2,000/39 ms TR/TE, 256×192 matrix size, r156uM resolution, 0.75 mm slice thickness, 0.25 mm slice gap, 40 × 30 cm FOV, 101 kHz bandwidth, and 4 NEX was used for acquired in coronal and axial geometries a multi-slice T2-weighted images. To reduce respiratory motion, the axial scan sequences were respiratory gated. All animal imaging, preparation, and maintenance was carried out in accordance with MD Anderson’s Institutional Animal Care and Use Committee policies and procedures.

IVIS-100 procedure has been described elsewhere (32).

### Surgical procedures

#### Orthotopic injection in kidney

First, 10^10 adeno-associated viral particles were resuspended in PBS (Thermo Fisher Scientific) and Matrigel matrix (Corning) 1:1 solution. Six- to nine-week-old mice were shaved and anesthetized using isoflurane (Henry Schein Animal Health). Analgesia was achieved with buprenorphine SR (0.1 mg/Kg BID) (Par Parmaceutical) via subcutaneous injection, and shaved skin was disinfected with 70% ethanol and betadine (Dynarex). A 1-cm incision was performed on the left flank through the skin/subcutaneous and muscular/peritoneal layers. Left kidney was exposed and 20 uL of viral resuspension was introduced by Hamilton syringe into the organ by a subcapsular injection. Hemostasis was controlled with a bipolar cautery (Bioseb) if needed. The kidney was carefully repositioned into the abdominal cavity, and muscular/peritoneal planes were closed individually by absorbable sutures. The skin/subcutaneous planes were closed using metal clips. Mice were monitored daily for the first three days, and twice/week thereafter for signs of tumor growth by manual palpation, bioluminescence imaging, and/or MRI when appropriate.

### Euthanasia, necropsy, and tissues collection

Mice were euthanized by exposure to CO_2_ followed by cervical dislocation. A necropsy form was filled in with mouse information, tumor size and weight, infiltrated organ annotations, and metastasis number and location.

### Tumor cell isolation and culture

*Ex vivo* cultures from primary tumor explants were generated by mechanical dissociation and incubation for 1 hour at 37°C with a solution of collagenase IV/dispase (2 mg/mL) (Invitrogen), resuspended in DMEM (Lonza) and filtered. Cells derived from tumor dissociation and digestion were plated on gelatin 0.1% (Millipore Sigma) coated plates and cultured in DMEM (Lonza) supplemented with 20% FBS (Lonza) and 1% penicillin–streptomycin, and kept in culture for less than 5 passages.

### Staining

Immunohistochemistry (IHC) and immunofluorescence (IF) were performed as previously described (32). *Antibodies list*: RFP (Thermo Fisher, cat. #MA5-15257), GFP (Abcam, cat. #13970), vimentin (Abcam, cat. #ab8978), Pax8 (Proteintech, cat. #10336-1-AP), Ki67 (Thermo Fisher, cat. #MA5-14520).

#### Multispectral imaging using the Vectra

Microwave treatment (MWT) was applied to perform antigen retrieval, quench endogenous peroxidases, and remove antibodies from earlier staining procedures. Akoya Biosciences AR6 antigen retrieval buffer (pH 6) was used for vimentin and RFP staining while Akoya Biosciences AR9 antigen retrieval buffer (pH 9) was used for Pax8 staining. The slides were stained with primary antibodies against RFP, Pax8, and vimentin, corresponding HRP conjugated secondary antibodies, and subsequently TSA dyes to generate Opal signal (vimentin, Opal 570; RFP, Opal 620; and Pax8, Opal 690). The slides were scanned with the Vectra 3 image scanning system (Caliper Life Sciences), and signals were unmixed and reconstructed into a composite image with Vectra inForm software 2.4.8.

*Estimation of purity* was calculated as percentage of positive area for Tdt IHC staining. Free online tool IHC profiler (33) was used for quantification.

### Metaphase spread and chromosome count

IF on metaphasic spread was obtained as previously described with few modifications (34). Cultures were treated with 100 ng/mL−1 nocodazole for 8 hours overnight, collected by trypsinization, resuspended in 0.2% (w/v) KCl and 0.2% (w/v) trisodium citrate hypotonic buffer at room temperature (20–22°C) for 5 to 10 min, and cytocentrifuged onto SuperFrost Plus glass slides (MenzelGlaser) at 450g for 10 min in a Shandon Cytospin 4. Slides were fixed at room temperature for 10 min in 1× PBS with 4% (v/v) formaldehyde, permeabilized for 10 min at room temperature in KCM buffer (120 mM KCl, 20 mM NaCl, 10 mM Tris (pH 7.5) and 0.1% (v/v) Triton X-100), and blocked with 5% goat serum PBS1X 0.1%Triton 100x BSA 3% for 30 min at room temperature. Slides were incubated with primary antibody diluted in antibody dilution buffer (PBS1X 0.1%Triton 100x BSA 3%) for 1 hr at room temperature, washed in 1× PBST (1× PBS with 0.1% (v/v)), incubated with secondary antibody diluted in antibody dilution buffer for 30 min at room temperature, washed with 1× PBST and stained for DNA with DAPI. Primary antibody: anti-centromere (ACA) (1:250; Antibodies Incorporated). Secondary antibody: goat anti-human conjugated to Alexa Fluor 488 (A-11013).

### Karyotype analysis

Exponentially growing cells in 10-cm petri dishes were treated with colcemid (0.04 ug/mL media) and incubated at 37°C for 2 to 3 hours. Cells were trypsinized and the single-cell suspension was collected in 15 mL conical centrifuge tubes. The cell suspension was centrifuged at 1500 rpm for 7 min. The supernatant was discarded, and the cell pellet was resuspended in 7 mL of hypotonic solution (0.075 M potassium chloride) and incubated for 20 min at room temperature. Then 2 mL of fixative (methanol and acetic acid, 3:1 v/v) was added to the cells, mixed, and centrifuged. The cells were washed thrice with fresh fixative and resuspended in about 1 mL of fixative. A few drops of cell suspension were placed on wet glass slides and allowed to air dry.

#### G banding

Slides were optimally aged (3 days at 65°C) and then Giemsa banded using trypsin EDTA solution following routine laboratory techniques. G-banded metaphase spreads were photographed using an 80i Nikon microscope and Applied Spectral Imaging karyotyping system (ASI, Vista, CA). A minimum of ten metaphases were karyotyped (35).

#### Spectral karyotyping (SKY)

Spectralkaryotyping was performed on these slides using the human SKY probe (Applied Spectral Imaging, Carlsbad, CA) according to the manufacturer’s protocol. Images were captured using 80i Nikon microscope and Applied Spectral Imaging (ASI) karyotyping system. A minimum of ten metaphases were karyotyped (36).

### Whole-exome sequencing of murine DNA

Illumina-compatible mouse exome libraries were prepared using the Agilent SureSelect protocol. Briefly, 1000 ng of Biorupter Ultrasonicator (Diagenode)–sheared, RNase-treated gDNA was used to construct sequencing libraries using the Agilent SureSelectXT Reagent Kit (Agilent Technologies). Libraries were prepared for capture with 6 cycles of PCR amplification, then assessed for size distribution on the 4200 TapeStation High Sensitivity D1000 ScreenTape (Agilent Technologies) and for quantity using the Qubit dsDNA HS Assay Kit (ThermoFisher). Exon target capture was performed using the Agilent SureSelectXT Mouse All Exon Kit. Following capture, index tags were added to the exon-enriched libraries using seven cycles of PCR. The indexed libraries were then assessed for size distribution using the Agilent TapeStation and quantified using the Qubit dsDNA HS Assay Kit respectively. Libraries were pooled, quantified by qPCR using the KAPA Library Quantification Kit (KAPABiosystems), and sequenced on the NovaSeq6000 using the 150 bp paired-end format.

### Next-generation sequencing of murine DNA

Exome libraries and whole genome-libraries were prepared and sequenced using a modified protocol originally described in Msaouel et al (37). Modifications to he protocol for murine exome sequencing were use of 1000ng of treated gDNA, performing only 6 cycles of PCR amplification, and usage of the Agilent SureSelectXT Mouse All Exon Kit for exon target capture. For murine whole - genome sequencing, after adapter ligation, libraries were only amplified by 2 cycles of PCR. Equimolar quantities of the whole-genome indexed libraries were multiplexed, with 18 libraries per pool. Results from 13 of the 18 libraries were used in our analysis. All pooled libraries were sequenced on an Illumina NovaSeq6000 using the 150 bp paired-end format.

### Bioinformatic processing of high-throughput sequencing data

The bioinformatic processing pipeline of raw whole-exome (WES) and whole-genome (WGS) high-throughput sequencing data was adapted for murine data from the protocol used in Seth et al (38). Reads were aligned to the mouse genome reference (mm10) using Burrows-Wheeler Aligner (BWA) with a seed length of 40 and a maximum edit distance of 3 (allowing for distance % 2 in the seed) (39). BAM files were further processed according to GATK Best Practices, including removal of duplicate reads, realignment around indels, and base recalibration (40,41).

### Analysis of sgRNA performance

Expected cut sites of sgRNAs were analyzed using CRISPResso2 (42). BAM files were first filtered with SAMtools (39) to contain reads spanning a 50-bp region centered around the expected sgRNA cut site and passed to CRISPResso2 in “CRISPRessoWGS” mode. The allele frequency of each base position around the cut site window was extracted from the CRISPResso2 results. An odds ratio for probability of a base position difference from the reference genome for each tumor sample and its respective matched normal sample was calculated by Fisher’s exact test by counting the number of base alterations observed at each cut site window position. The odds ratios were transformed by natural log and z-transformation against the average log-odds ratio for all base positions of the same gene (43). The z-transformed log-odds ratios were then averaged across all gene cut sites for a sample to summarize the overall editing efficiency of the sgRNAs delivered to each mouse (44). Genes were considered altered if at least 3 reads with the same pattern of base alteration were detected at the expected sgRNA cut site.

### Identification of somatic copy number profiles and events

CNVkit (45) was used to derive somatic copy number profiles from WES data using a panel of normal samples consisting of all the matched normal samples across all mice sequenced in this study. The targeted exome bed file for the Agilent SureSelect All Mouse Exon V1 was downloaded from Agilent with the original mm9 coordinates and was then converted to mm10 using CrossMap v0.3.4 (46) for use by CNVkit. Occurrence of CNVs in focal regions of the genome (e.g., C11qE, Cdkn2a/b) were called if all exons spanning the region of interest had a absolute weighted average log_2_ read-depth ratio of ≥0.4. Otherwise, GISTIC2 was run with amplification and deletion thresholds of 0.2, using gene-level assumptions for significance, along with additional broad-level analysis. The GISTIC2 reference genome file for mm10 was acquired, and no marker file was necessary (47), (48).

Sequenza (49) was used to derive somatic copy number profiles from WGS data using each sample’s matched normal sample. A modified version of the “copynumber” R package (https://github.com/aroneklund/copynumber) was used to create the necessary support objects and allowed for usage of the mm10 genome in the deployment of Sequenza on the mouse BAM files. To assign ploidy to WGS samples, purity was first estimated by Ki67 fluorescence as previously described, and the ploidy with the largest predicted probability at the estimated purity was selected from the Sequenza cellularity-ploidy prediction table.

### Construction of tumor progression sample tree representation

The sample progression tree representation of tumors was first constructed with hierarchical clustering using the complete linkage algorithm (50) and the hamming distance between samples. The hamming distance was calculated as the number of non-driver somatic mutations (as determined from WES) shared by any two samples as a fraction of the total number of non-somatic mutations contained by either sample. Visualizations of sample progression trees were manually generated. Branch lengths of 0 were collapsed to its direct ancestor node. Only mutations detected in all descendants of a branch were considered.

### Statistical analysis of clinical RCC cohort data

Processed clinical, copy number, somatic mutation, and molecular characterization data from The Cancer Genome Atlas’ (TCGA) pan-kidney (KIPAN) tumor sample cohort were obtained from Ricketts et al (51). TCGA profiling data was then augmented with arm-level copy number calls, aneuploidy score, and whole-genome doubling (WGD) status as determined by Taylor et al (16). The aneuploidy score was then transformed to calculate a fraction of genome altered (fCNA) as described in Taylor et al (16). Only samples with copy number data and aneuploidy data were included in the analysis of TCGA pan-kidney cohort. TCGA tumors with sarcomatoid features were manually annotated as described in Bokouny et al (52). Clinical data used for confirmation of genomic effects of 9p loss on WGD and aneuploidy were acquired from the TRACERx renal cell cancer cohort and a renal cell carcinoma cohort from Memorial Sloan Kettering Cancer Center kidney cancer cohort (MSKCC, unpublished) (10,16,51). The aneuploidy score for TRACERx samples was calculated using the arm-level chromosome alteration calls from TRACERx directly and then converted to an fCNA value as described in Taylor et al (16).

### Summary of methods for uRCC MSK cohort

RCC tumor specimens from 134 patients were procured from the Memorial Sloan Kettering (MSK) Pathology Department after Ethics Review Board approval. Primary and metastatic deposit specimens were reviewed by a specialized genitourinary pathologist (Y.B.C) and a diagnosis of high-grade unclassified renal cell carcinoma was made. Clinicopathologic and molecular data for 62 of these patients have been reported in a previous publication (8).

Macro-dissected tumor and paired adjacent normal kidney tissue or blood were sent for DNA extraction and sequencing at the Integrated Genomic Operations Core of MSK or Molecular Diagnostics Service laboratory of Department of Pathology. Sequencing was done on both the tumor and matched-normal samples using the MSK-IMPACT gene panel (MSK-IMPACT^®^) (53). Samples were sequenced at an average depth of 500x.

Raw sequencing data was aligned to a reference genome (b37) and somatic variants were called using a previously validated pipeline. Briefly, four different variant calling tools were used for this purpose: MuTect2 (part of GATKv4.1.4.1 (40)), Strelka2 v2.9.10 (54), Varscan v2.4.3 (55) and Platypus (56). Ancillary filters were then applied to obtain high-accuracy mutations, these included: a coverage of at least 10 reads in the tumor, with 5 or more supporting the variant of interest, a variant allele frequency (vAF) ≥5% in the tumor, and a vAF<7% in the matched normal sample. Only somatic nonsynonymous exonic mutations were considered, and single-nucleotide variants (SNVs) identified at a frequency >1% in dbSNP (57) or 1000Genomes project (57) were removed. All variant calls were manually reviewed by investigators (R.G.D) for additional accuracy.

Allele-specific copy number analysis (ASCN) and purity estimation were done using the FACETS algorithm v0.5.6. (58). Finally, inference of arm-level and genome-doubling events was performed using a public R package (https://github.com/mskcc/facets-suite). All copy-number variations (CNVs) in autosomal chromosomes were considered, regardless of length.

Informed consent was obtained after the nature and possible consequences of the studies were explained.

### B-allele frequency comparison

Murine B-allele frequencies were calculated using the snp-pileup script from the FACETS software package on WGS samples (58). The VCF of identified murine SNP locations was obtained from the Wellcome Sanger Institute, Mouse Genome Project version 5, dbSNP142 (59,60). The snp-pileup counts were then utilized to determine the allele frequency of these common murine SNPs. Heterozygous SNPs were identified if the B-allele (alternative nucleotide) frequency (BAF) was 0.2 < BAF < 0.8 with minimum coverage of 15x in the normal tumor sample. B-allele frequencies of heterozygous SNPs identified in each mouse’s normal tissue sample was plotted against corresponding tissue sample BAFs for the same SNP.

### Statistical analysis

Data are presented as the mean ± s.d and percentages. Comparisons between biological replicates were performed using a two-tailed Student’s *t* test or Mann-Whitney U test. Results from survival experiments were analyzed with a log-rank (Mantel–Cox) test and expressed as Kaplan–Meier survival curves. Results from contingency tables were analyzed using the two-tailed Fisher’s exact test or chi-test for multiple comparisons. (GraphPad software). Group size was determined on the basis of the results of preliminary experiments. No statistical methods were used to determine sample size. Group allocation and analysis of outcome were not performed in a blinded manner. Samples that did not meet proper experimental conditions were excluded from the analysis.

## Acknowledgments

We wish to thank Drs. Jintan Liu and Mike Peoples for discussions and advices, Drs. Joshua Goldenberg and Stefania Napolitano for technical support, and all members of the Draetta lab for discussions and reagents. We would particularly like to thank Shan Jiang, and the MDACC Department of Veterinary Medicine, for valuable support in animals handling, Dr. Charles Kingsley, Jorge Delacerda, and the MDACC Small Animals Imaging Facility for their constant willingness. We wish to thank Dr. Multani Asha and the MDACC Cytogenetics and Cell Authentication Core; Dr. Kazuhiro Oka and the Baylor Gene Vector Core. We thank Genscript for support and service. We wish to acknowledge the Advanced Technology Genomics Core (Erika Thompson, Dr. Hongli Tang, Steven Bates) at MDACC and the CA016672(ATGC) Core Grant. The authors acknowledge the support of the High Performance Computing for research facility at the University of Texas MD Anderson Cancer Center for providing computational resources that have contributed to the research results reported in this paper. Special thanks to Sarah Townsend for reviewing the entire manuscript. We thank Viviana Gradinaru and Benjamin Deverman for AAV PHP.eB (Catalog # 28304-PHPeB).

## Author contributions

Conceptualization, F.C., G.G. A.C., K.S., A.S., J.A.K., J.G., E.J., A.V., G.F.D., Z.B., E.M.V.A., T.C., Y.C., R.G D., A.A H., K.C.A., A.F., T.P.H., G.G.M.,P.M., K.L., S.T., C.A.B., N.M.T.; Methodology, F.C., J.K.H., C.A.B., A.C., G.G.; Software, X.S., J.Z., J.K.H., C.A.B.; Formal Analysis, F.C., J.K.H., L.P.; Investigation, F.C., J.K.H., L.P., E.D.P., T.G., H.T., M.S., T.N.L., R.X., C.Z.; Access to patient samples: P.M., N.M.T., J.K.; Writing – Original Draft, F.C., J.K.H. and G.G.; Writing – Review & Editing, F.C., J.K.H., G.G., L.P., A.C., E.D.P., A.V., N.M.T., P.M., C.A.B., Z.B.; Visualization, F.C., J.K.H., L.P., G.G. ; Supervision, G.G., C.A.B., A.C., and N.M.T.; Funding Acquisition, F.C., A.C., N.M.T., G.F.D., and G.G.

## Data and materials availability

All data is available in the manuscript or the supplementary materials.

## References

1. Linehan WM, Ricketts CJ. RCC — advances in targeted therapeutics and genomics. Nature Reviews Urology 2017;14(2):76–8 doi 10.1038/nrurol.2016.260.

2. Lindgren D, Sjolund J, Axelson H. Tracing Renal Cell Carcinomas back to the Nephron. Trends Cancer 2018;4(7):472–84 doi 10.1016/j.trecan.2018.05.003.

3. Wolf MM, Kimryn Rathmell W, Beckermann KE. Modeling clear cell renal cell carcinoma and therapeutic implications. Oncogene 2020;39(17):3413–26 doi 10.1038/s41388-020-1234-3.

4. Gu YF, Cohn S, Christie A, McKenzie T, Wolff N, Do QN, et al. Modeling Renal Cell Carcinoma in Mice: Bap1 and Pbrm1 Inactivation Drive Tumor Grade. Cancer Discov 2017;7(8):900–17 doi 10.1158/2159-8290.CD-17-0292.

5. Nargund AM, Pham CG, Dong Y, Wang PI, Osmangeyoglu HU, Xie Y, et al. The SWI/SNF Protein PBRM1 Restrains VHL-Loss-Driven Clear Cell Renal Cell Carcinoma. Cell Rep 2017;18(12):2893–906 doi 10.1016/j.celrep.2017.02.074.

6. Malouf GG, Ali SM, Wang K, Balasubramanian S, Ross JS, Miller VA, et al. Genomic Characterization of Renal Cell Carcinoma with Sarcomatoid Dedifferentiation Pinpoints Recurrent Genomic Alterations. Eur Urol 2016;70(2):348–57 doi 10.1016/j.eururo.2016.01.051.

7. Wang Z, Kim TB, Peng B, Karam J, Creighton C, Joon A, et al. Sarcomatoid Renal Cell Carcinoma Has a Distinct Molecular Pathogenesis, Driver Mutation Profile, and Transcriptional Landscape. Clin Cancer Res 2017;23(21):6686–96 doi 10.1158/1078-0432.CCR-17-1057.

8. Chen YB, Xu J, Skanderup AJ, Dong Y, Brannon AR, Wang L, et al. Molecular analysis of aggressive renal cell carcinoma with unclassified histology reveals distinct subsets. Nat Commun 2016;7:13131 doi 10.1038/ncomms13131.

9. Keskin SK, Msaouel P, Hess KR, Yu KJ, Matin SF, Sircar K, et al. Outcomes of Patients with Renal Cell Carcinoma and Sarcomatoid Dedifferentiation Treated with Nephrectomy and Systemic Therapies: Comparison between the Cytokine and Targeted Therapy Eras. J Urol 2017;198(3):530–7 doi 10.1016/j.juro.2017.04.067.

10. Turajlic S, Xu H, Litchfield K, Rowan A, Chambers T, Lopez JI, et al. Tracking Cancer Evolution Reveals Constrained Routes to Metastases: TRACERx Renal. Cell 2018;173(3):581–94 e12 doi 10.1016/j.cell.2018.03.057.

11. Ricketts CJ, De Cubas AA, Fan H, Smith CC, Lang M, Reznik E, et al. The Cancer Genome Atlas Comprehensive Molecular Characterization of Renal Cell Carcinoma. Cell Rep 2018;23(12):3698 doi 10.1016/j.celrep.2018.06.032.

12. Weber J, Ollinger R, Friedrich M, Ehmer U, Barenboim M, Steiger K, et al. CRISPR/Cas9 somatic multiplex-mutagenesis for high-throughput functional cancer genomics in mice. Proc Natl Acad Sci U S A 2015;112(45):13982–7 doi 10.1073/pnas.1512392112.

13. Wang G, Chow RD, Ye L, Guzman CD, Dai X, Dong MB, et al. Mapping a functional cancer genome atlas of tumor suppressors in mouse liver using AAV-CRISPR-mediated direct in vivo screening. Sci Adv 2018;4(2):eaao5508 doi 10.1126/sciadv.aao5508.

14. Xue W, Chen S, Yin H, Tammela T, Papagiannakopoulos T, Joshi NS, et al. CRISPR-mediated direct mutation of cancer genes in the mouse liver. Nature 2014;514(7522):380–4 doi 10.1038/nature13589.

15. Davis A, Gao R, Navin N. Tumor evolution: Linear, branching, neutral or punctuated? Biochim Biophys Acta Rev Cancer 2017;1867(2):151–61 doi 10.1016/j.bbcan.2017.01.003.

16. Taylor AM, Shih J, Ha G, Gao GF, Zhang X, Berger AC, et al. Genomic and Functional Approaches to Understanding Cancer Aneuploidy. Cancer Cell 2018;33(4):676–89 e3 doi 10.1016/j.ccell.2018.03.007.

17. Sansregret L, Vanhaesebroeck B, Swanton C. Determinants and clinical implications of chromosomal instability in cancer. Nat Rev Clin Oncol 2018;15(3):139–50 doi 10.1038/nrclinonc.2017.198.

18. Cancer Genome Atlas Research N, Linehan WM, Spellman PT, Ricketts CJ, Creighton CJ, Fei SS, et al. Comprehensive Molecular Characterization of Papillary Renal-Cell Carcinoma. N Engl J Med 2016;374(2):135–45 doi 10.1056/NEJMoa1505917.

19. Chen F, Zhang Y, Senbabaoglu Y, Ciriello G, Yang L, Reznik E, et al. Multilevel Genomics-Based Taxonomy of Renal Cell Carcinoma. Cell Rep 2016;14(10):2476–89 doi 10.1016/j.celrep.2016.02.024.

20. Braun DA, Hou Y, Bakouny Z, Ficial M, Sant’ Angelo M, Forman J, et al. Interplay of somatic alterations and immune infiltration modulates response to PD-1 blockade in advanced clear cell renal cell carcinoma. Nature Medicine 2020;26(6):909–18 doi 10.1038/s41591-020-0839-y.

21. Hou W, Ji Z. Generation of autochthonous mouse models of clear cell renal cell carcinoma: mouse models of renal cell carcinoma. Exp Mol Med 2018;50(4):30 doi 10.1038/s12276-018-0059-4.

22. Sobczuk P, Brodziak A, Khan MI, Chhabra S, Fiedorowicz M, Wełniak-Kamińska M, et al. Choosing The Right Animal Model for Renal Cancer Research. Transl Oncol 2020;13(3):100745 doi 10.1016/j.tranon.2020.100745.

23. Gao R, Davis A, McDonald TO, Sei E, Shi X, Wang Y, et al. Punctuated copy number evolution and clonal stasis in triple-negative breast cancer. Nat Genet 2016;48(10):1119–30 doi 10.1038/ng.3641.

24. Bouchard M, Souabni A, Mandler M, Neubüser A, Busslinger M. Nephric lineage specification by Pax2 and Pax8. Genes Dev 2002;16(22):2958–70 doi 10.1101/gad.240102.

25. Chiou S-H, Winters IP, Wang J, Naranjo S, Dudgeon C, Tamburini FB, et al. Pancreatic cancer modeling using retrograde viral vector delivery and in vivo CRISPR/Cas9-mediated somatic genome editing. Genes Dev 2015;29(14):1576–85 doi 10.1101/gad.264861.115.

26. Madisen L, Zwingman TA, Sunkin SM, Oh SW, Zariwala HA, Gu H, et al. A robust and high-throughput Cre reporting and characterization system for the whole mouse brain. Nat Neurosci 2010;13(1):133–40 doi 10.1038/nn.2467.

27. Madisen L, Garner AR, Shimaoka D, Chuong AS, Klapoetke NC, Li L, et al. Transgenic mice for intersectional targeting of neural sensors and effectors with high specificity and performance. Neuron 2015;85(5):942–58 doi 10.1016/j.neuron.2015.02.022.

28. Safran M, Kim WY, Kung AL, Horner JW, DePinho RA, Kaelin WG, Jr. Mouse reporter strain for noninvasive bioluminescent imaging of cells that have undergone Cre-mediated recombination. Mol Imaging 2003;2(4):297–302 doi 10.1162/153535003322750637.

29. Schwarz LA, Miyamichi K, Gao XJ, Beier KT, Weissbourd B, DeLoach KE, et al. Viral-genetic tracing of the input-output organization of a central noradrenaline circuit. Nature 2015;524(7563):88–92 doi 10.1038/nature14600.

30. Simpson EM, Korecki AJ, Fornes O, McGill TJ, Cueva-Vargas JL, Agostinone J, et al. New MiniPromoter Ple345 (NEFL) Drives Strong and Specific Expression in Retinal Ganglion Cells of Mouse and Primate Retina. Hum Gene Ther 2019;30(3):257–72 doi 10.1089/hum.2018.118.

31. Chan KY, Jang MJ, Yoo BB, Greenbaum A, Ravi N, Wu W-L, et al. Engineered AAVs for efficient noninvasive gene delivery to the central and peripheral nervous systems. Nat Neurosci 2017;20(8):1172–9 doi 10.1038/nn.4593.

32. Genovese G, Carugo A, Tepper J, Robinson FS, Li L, Svelto M, et al. Synthetic vulnerabilities of mesenchymal subpopulations in pancreatic cancer. Nature 2017;542(7641):362–6 doi 10.1038/nature21064.

33. Varghese F, Bukhari AB, Malhotra R, De A. IHC Profiler: an open source plugin for the quantitative evaluation and automated scoring of immunohistochemistry images of human tissue samples. PLoS One 2014;9(5):e96801–e doi 10.1371/journal.pone.0096801.

34. Cesare AJ, Kaul Z, Cohen SB, Napier CE, Pickett HA, Neumann AA, et al. Spontaneous occurrence of telomeric DNA damage response in the absence of chromosome fusions. Nature Structural & Molecular Biology 2009;16(12):1244–51 doi 10.1038/nsmb.1725.

35. Wang R, Chu GCY, Wang X, Wu JB, Hu P, Multani AS, et al. Establishment and characterization of a prostate cancer cell line from a prostatectomy specimen for the study of cellular interaction. International Journal of Cancer 2019;145(8):2249–59 doi 10.1002/ijc.32370.

36. Deriano L, Chaumeil J, Coussens M, Multani A, Chou Y, Alekseyenko AV, et al. The RAG2 C terminus suppresses genomic instability and lymphomagenesis. Nature 2011;471(7336):119–23 doi 10.1038/nature09755.

37. Msaouel P, Malouf GG, Su X, Yao H, Tripathi DN, Soeung M, et al. Comprehensive Molecular Characterization Identifies Distinct Genomic and Immune Hallmarks of Renal Medullary Carcinoma. Cancer Cell 2020;37(5):720–34.e13 doi 10.1016/j.ccell.2020.04.002.

38. Seth S, Li C-Y, Ho IL, Corti D, Loponte S, Sapio L, et al. Pre-existing Functional Heterogeneity of Tumorigenic Compartment as the Origin of Chemoresistance in Pancreatic Tumors. Cell Reports 2019;26(6):1518–32.e9 doi https://doi.org/10.1016/j.celrep.2019.01.048.

39. Li H, Durbin R. Fast and accurate short read alignment with Burrows-Wheeler transform. Bioinformatics 2009;25(14):1754–60 doi 10.1093/bioinformatics/btp324.

40. DePristo MA, Banks E, Poplin R, Garimella KV, Maguire JR, Hartl C, et al. A framework for variation discovery and genotyping using next-generation DNA sequencing data. Nature Genetics 2011;43(5):491–8 doi 10.1038/ng.806.

41. Van der Auwera GA, Carneiro MO, Hartl C, Poplin R, Del Angel G, Levy-Moonshine A, et al. From FastQ data to high confidence variant calls: the Genome Analysis Toolkit best practices pipeline. Curr Protoc Bioinformatics 2013;43(1110):11.0.1-.0.33 doi 10.1002/0471250953.bi1110s43.

42. Clement K, Rees H, Canver MC, Gehrke JM, Farouni R, Hsu JY, et al. CRISPResso2 provides accurate and rapid genome editing sequence analysis. Nature Biotechnology 2019;37(3):224–6 doi 10.1038/s41587-019-0032-3.

43. Xue W, Chen S, Yin H, Tammela T, Papagiannakopoulos T, Joshi NS, et al. CRISPR-mediated direct mutation of cancer genes in the mouse liver. Nature 2014;514(7522):380–4 doi 10.1038/nature13589.

44. Mueller S, Engleitner T, Maresch R, Zukowska M, Lange S, Kaltenbacher T, et al. Evolutionary routes and KRAS dosage define pancreatic cancer phenotypes. Nature 2018;554(7690):62–8 doi 10.1038/nature25459.

45. Talevich E, Shain AH, Botton T, Bastian BC. CNVkit: Genome-Wide Copy Number Detection and Visualization from Targeted DNA Sequencing. PLoS Comput Biol 2016;12(4):e1004873–e doi 10.1371/journal.pcbi.1004873.

46. Zhao H, Sun Z, Wang J, Huang H, Kocher J-P, Wang L. CrossMap: a versatile tool for coordinate conversion between genome assemblies. Bioinformatics 2013;30(7):1006–7 doi 10.1093/bioinformatics/btt730.

47. Mermel CH, Schumacher SE, Hill B, Meyerson ML, Beroukhim R, Getz G. GISTIC2.0 facilitates sensitive and confident localization of the targets of focal somatic copy-number alteration in human cancers. Genome Biology 2011;12(4):R41 doi 10.1186/gb-2011-12-4-r41.

48. Lange S, Engleitner T, Mueller S, Maresch R, Zwiebel M, González-Silva L, et al. Analysis pipelines for cancer genome sequencing in mice. Nature Protocols 2020;15(2):266–315 doi 10.1038/s41596-019-0234-7.

49. Favero F, Joshi T, Marquard AM, Birkbak NJ, Krzystanek M, Li Q, et al. Sequenza: allele-specific copy number and mutation profiles from tumor sequencing data. Annals of Oncology 2015;26(1):64–70 doi 10.1093/annonc/mdu479.

50. Müllner D. 12 September 2011.

51. Ricketts CJ, De Cubas AA, Fan H, Smith CC, Lang M, Reznik E, et al. The Cancer Genome Atlas Comprehensive Molecular Characterization of Renal Cell Carcinoma. Cell Reports 2018;23(1):313–26.e5 doi 10.1016/j.celrep.2018.03.075.

52. Bakouny Z, Braun DA, Shukla SA, Pan W, Gao X, Hou Y, et al. Integrative Molecular Characterization of Sarcomatoid and Rhabdoid Renal Cell Carcinoma Reveals Determinants of Poor Prognosis and Response to Immune Checkpoint Inhibitors. bioRxiv 2020:2020.05.28.121806 doi 10.1101/2020.05.28.121806.

53. Cheng DT, Mitchell TN, Zehir A, Shah RH, Benayed R, Syed A, et al. Memorial Sloan Kettering-Integrated Mutation Profiling of Actionable Cancer Targets (MSK-IMPACT): A Hybridization Capture-Based Next-Generation Sequencing Clinical Assay for Solid Tumor Molecular Oncology. J Mol Diagn 2015;17(3):251–64 doi 10.1016/j.jmoldx.2014.12.006.

54. Kim S, Scheffler K, Halpern AL, Bekritsky MA, Noh E, Källberg M, et al. Strelka2: Fast and accurate variant calling for clinical sequencing applications. bioRxiv 2017:192872 doi 10.1101/192872.

55. Koboldt DC, Zhang Q, Larson DE, Shen D, McLellan MD, Lin L, et al. VarScan 2: somatic mutation and copy number alteration discovery in cancer by exome sequencing. Genome Res 2012;22(3):568–76 doi 10.1101/gr.129684.111.

56. Rimmer A, Phan H, Mathieson I, Iqbal Z, Twigg SRF, Wilkie AOM, et al. Integrating mapping-, assembly- and haplotype-based approaches for calling variants in clinical sequencing applications. Nat Genet 2014;46(8):912–8 doi 10.1038/ng.3036.

57. Durbin RM, Altshuler D, Durbin RM, Abecasis GR, Bentley DR, Chakravarti A, et al. A map of human genome variation from population-scale sequencing. Nature 2010;467(7319):1061–73 doi 10.1038/nature09534.

58. Shen R, Seshan VE. FACETS: allele-specific copy number and clonal heterogeneity analysis tool for high-throughput DNA sequencing. Nucleic Acids Research 2016;44(16):e131–e doi 10.1093/nar/gkw520.

59. Keane TM, Goodstadt L, Danecek P, White MA, Wong K, Yalcin B, et al. Mouse genomic variation and its effect on phenotypes and gene regulation. Nature 2011;477(7364):289–94 doi 10.1038/nature10413.

60. Yalcin B, Wong K, Agam A, Goodson M, Keane TM, Gan X, et al. Sequence-based characterization of structural variation in the mouse genome. Nature 2011;477(7364):326–9 doi 10.1038/nature10432.

61. Lee J, Hong W-y, Cho M, Sim M, Lee D, Ko Y, et al. Synteny Portal: a web-based application portal for synteny block analysis. Nucleic Acids Research 2016;44(W1):W35–W40 doi 10.1093/nar/gkw310.

