## Supplementary Figures and Legends for "9p21 Loss Defines the Evolutionary Patterns of Aggressive Renal Cell Carcinomas"

**Supplementary Fig. S1.**

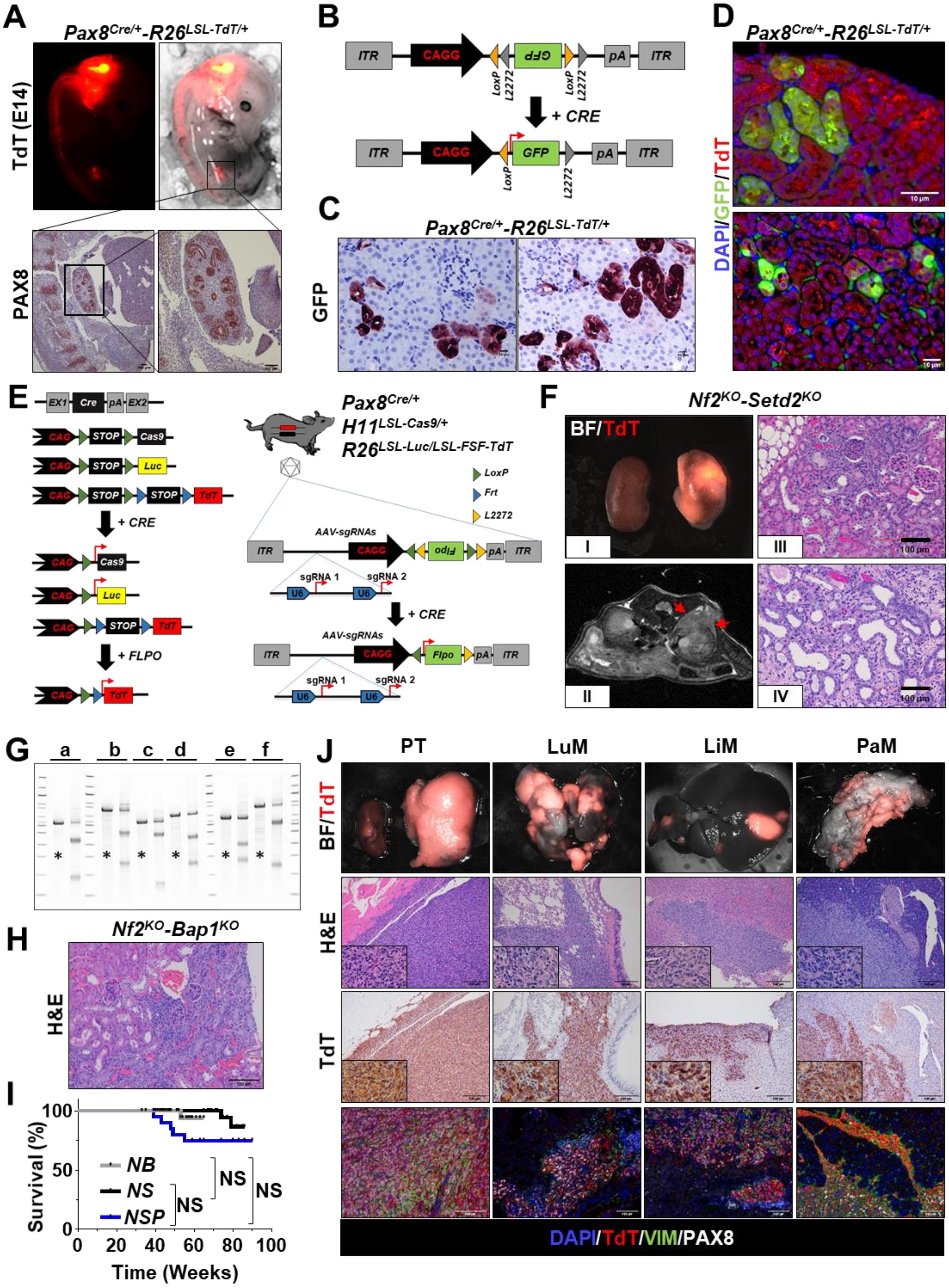

**A)** Representative E14 (embryonic day 14) *Pax8*$^{Cre/+}$-*R26*$^{LSL\text{-}TdT/+}$ embryos. The activation of the fluorescent reporter TdTomato (TdT) can be readily appreciated in the developing hindbrain, notochord, and kidney by both gross macroscopic and by microscopic immune histochemical analysis for Pax8. **B)** Schematic showing the AAV-based tracing system carrying a FLEx GFP reported sequence. Upon orthotopic delivery of $10^{10}$ infectious units into *Pax8*$^{Cre/+}$-*R26*$^{LSL\text{-}TdT/+}$ mice, the tissue-specific Cre recombinase drives the activation of the GFP reporter in the epithelial compartment of the nephron. **C-D)** IHC and IF analysis on representative FFPE sections stained with an antibody specific for GFP (c) and GFP and TdT (d). Tissues were harvested 10 days after orthotopic transduction with$10^{10}$ infectious units. Mosaic activation of the reporter in transduced epithelial cells of the nephron can be readily appreciated. **E)** Schematic showing the SM-GEMM design. The *Pax8*$^{Cre}$ strain was crossed with the conditional *H11*$^{LSL\text{-}Cas9}$, *R26*$^{LSL\text{-}LUC}$, and *R26*$^{LSL\text{-}FSF\text{-}TdT}$ strains to generate the *PCLT* model, allowing for tissue-specific activation of Cas9 and luciferase (Luc) in the renal epithelium and priming the system for the somatic, FlpO-mediated activation of the TdTomato (TdT) fluorescence reporter. Cancer-specific loss-of-function mutations are introduced via intraparenchymal delivery of adeno-associated viral (AAV) particles carrying specific sgRNA combinations. The latent TdT fluorescence reporter allows for the tracing of renal epithelial cells successfully transduced with AAV particles. The reporter is activated through a two-step recombination system upon Cre-mediated ablation of a LoxP-Stop-LoxP (LSL) cassette in the epithelial compartment of the kidney and FlpO-mediated mosaic removal of a Frt-Stop-Frt (FSF) cassette in AAV-targeted kidney epithelial cells. The latent Cre-inducible FlpO cassette carried by the AAV vector is activated via FLEx technology. Tight regulation of the TdT reporter allows for precise labeling of CRISPR-edited somatic cells. **F)** Pathological characterization of murine RCC obtained through somatic mosaic knockout of *Nf2* and *Setd2*. Mice developed low-grade tumors rarely presenting aggressive features. i) Gross specimens harvested 8 months post-transduction, ii) axial T2 MRI scan displaying a small cortical lesion 8 months post transduction, and iii-iv) hematoxylin-eosin (H&E) stained section from well-differentiated tumors harvested at 6 and 8 months post-transduction, respectively. Areas with papillary and tubular features can be readily appreciated. **G)** T7-endonuclease assay on MCT cells infected with CRISPR-V2 vector carrying a sgRNA targeting *Trp53* (a), *Nf2* (b), *Bap1* (c), *Setd2* (d), *Cdkn2a* (e), *Cdkn2b* (f). DNA of control vector–infected MCT cells were used as negative controls (*). **H)** Pathological characterization of murine RCC obtained through somatic mosaic knockout of *Nf2* and *Bap1*. **I)** Kaplan–Meier analysis of cancer-specific survival of *Nf2*$^{KO}$-driven murine tumors. Non-significant differences in survival are appreciated. NB: *Nf2*$^{KO}$-*Bap1*$^{KO}$ (N = 40); NS: *Nf2*$^{KO}$-*Setd2*$^{KO}$ (N = 20); NSP: *Nf2*$^{KO}$-*Setd2*$^{KO}$-*Trp53*$^{KO}$ (N = 24). **J)** Characterization of *Nf2*$^{KO}$-driven murine tumors upon genetic targeting of the murine locus syntenic to human 9p21.3 ($4q^{9p21}$): representative macroscopic images (*top panels*), hematoxylin/eosin (H&E), IHC, and IF analysis of specific lineage and tumor markers are reported. PT: primary tumor, LuM: lung metastasis, LiM: liver metastasis, PaM: pancreatic metastasis. N.S.: not significant by log-rank (Mantel–Cox) test.

**Supplementary Fig. S2.**

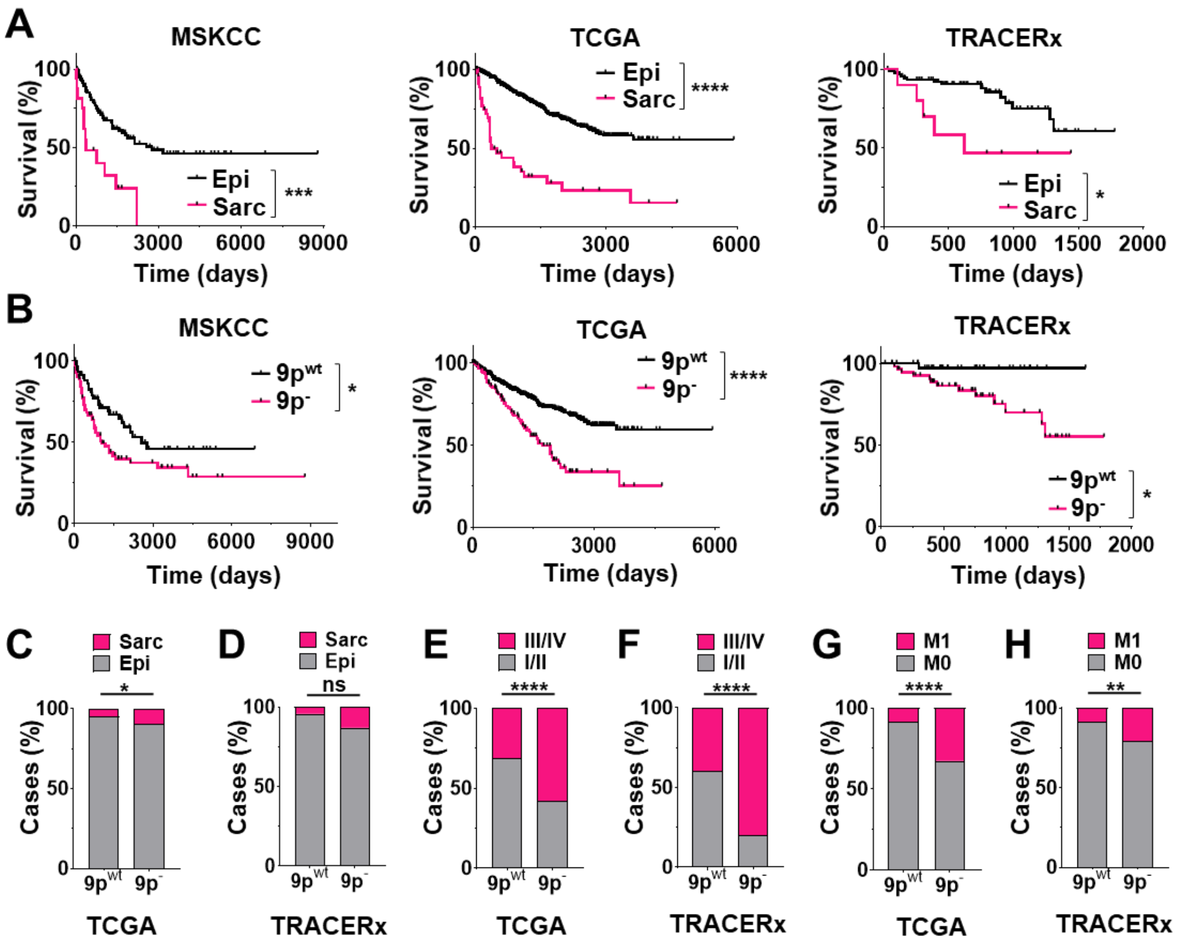


**A)** Kaplan–Meier survival analysis of human RCCs with and without sarcomatoid features in MSKCC (N = 16 vs N = 97) (*left panel*), TCGA (N = 45 vs N = 743) (*middle panel*) and TRACERx (N = 10 vs N = 91) (*right panel*) cohorts. **B)** Kaplan–Meier survival analysis of human RCCs with and without 9p loss features in MSKCC (N = 72 vs N = 62) (*left panel*), TCGA (N = 140 vs N = 658) (*middle panel*) and TRACERx (N = 57 vs N = 38) (*right panel*) cohort. **C-D)** Bar chart showing the prevalence of sarcomatoid features in $9p^{wt}$ and $9p^{-}$ cases in TCGA pan-RCC dataset (N = 648 vs N = 140) (C) and TRACERx RCC dataset (N = 45 vs N = 61) (D). **E-F)** Bar chart showing the prevalence of stage I/II and stage III/IV features in $9p^{wt}$ and $9p^{-}$ cases in TCGA pan-RCC dataset (N = 628 vs N = 136) (E) and TRACERx RCC dataset (N = 45 vs N = 61) (F). **G-H)** Bar chart showing the prevalence of metastasis features in $9p^{wt}$ and $9p^{-}$ cases in TCGA pan-RCC dataset (N = 628 vs N = 136) (G) and TRACERx RCC dataset (N = 45 vs N = 61) (H). N.S.: not significant, * $P < 0.05$, ** $P < 0.01$, *** $P < 0.001$, **** $P < 0.0001$ by Fisher's exact test (C-H), log-rank (Mantel–Cox) test (A,B).

**Supplementary Fig. S3.**

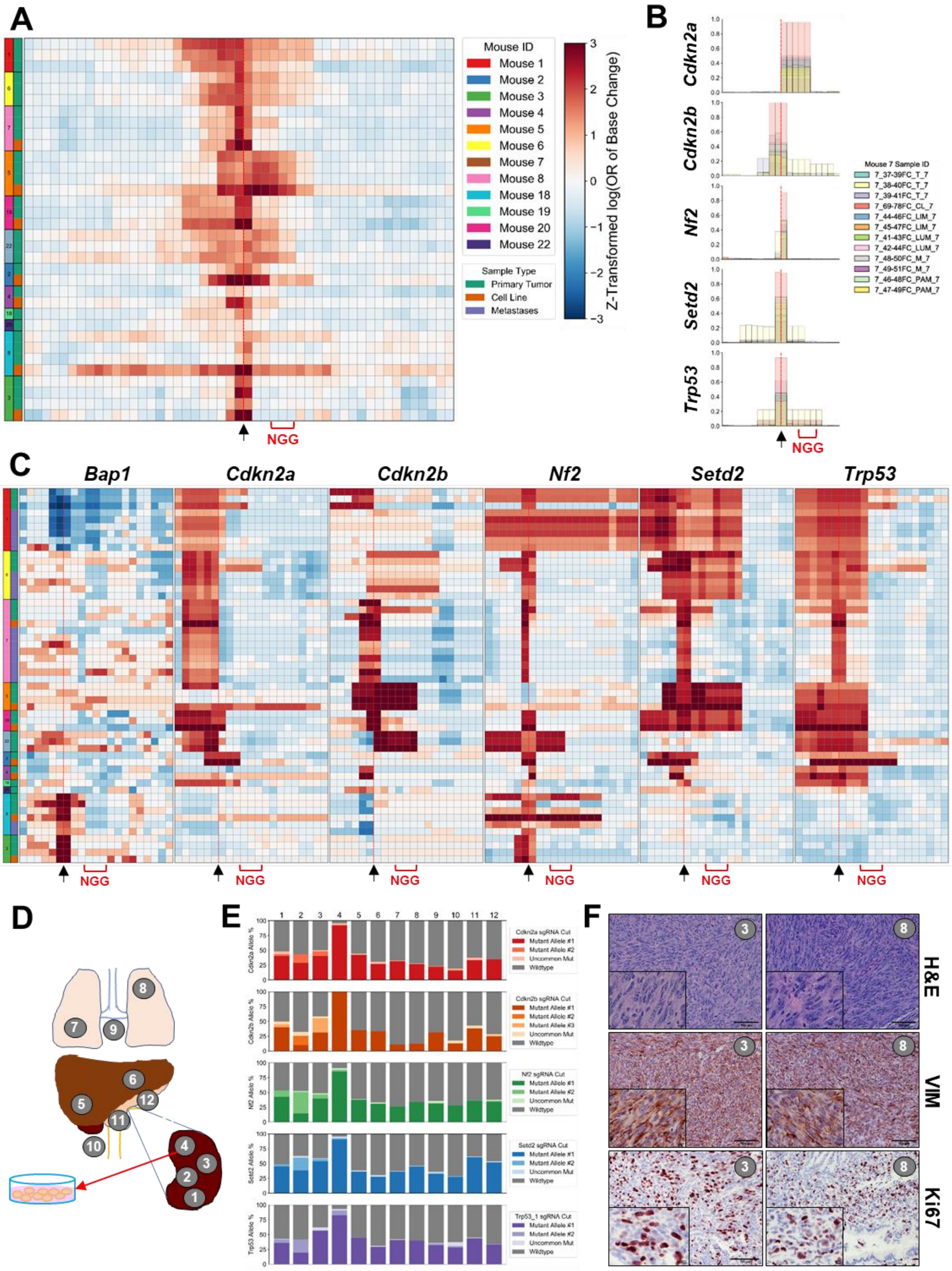

**A)** Heatmap showing the average Z-transformed log odds ratio across all edited genes for the likelihood of specific base alterations in any reads spanning an expected cut site. Data in figure were generated from WES analysis of primary tumor samples (N = 34 samples across 12 mice). **B)** Overlapping bar chart of the frequency of edited bases at each expected cut site for each gene engineered in all multiregional (primary and metastasis) samples for a single representative mouse M#7. **C)** Heatmaps showing the individual Z-transformed log odds ratios of base alteration at individual cut sites for each gene targeted by the reported sgRNAs. Primary and metastatic sites were profiled by WES (N = 54 samples across 12 mice). Arrows represent predicted cut site of sgRNA upstream of the PAM sequence (NGG) (A-C). **D)** Representative schematic of multiregional sampling (mouse M#7).1, 2, 3: primary tumor, 4: cell line established from primary site, 5, 6: liver metastasis, 7, 8: lung metastasis, 9: diaphragmatic metastasis, 10: mesenteric metastasis, 11, 12: pancreatic metastasis. **E)** Stacked bar charts highlighting the most abundant alterations introduced by CRISPR-Cas9 system in each targeted gene across all samples from the representative mouse in D. The prevalence of a few events across the samples is readily appreciated. **F)** Representative H&E (*upper panel*) and IHC staining for vimentin (VIM) and Ki67 (*middle and lower panel*, respectively) of histopathological sections obtained from a primary tumor and related lung metastasis area, showing highly consistent morphometric features.

**Supplementary Fig. S4.**

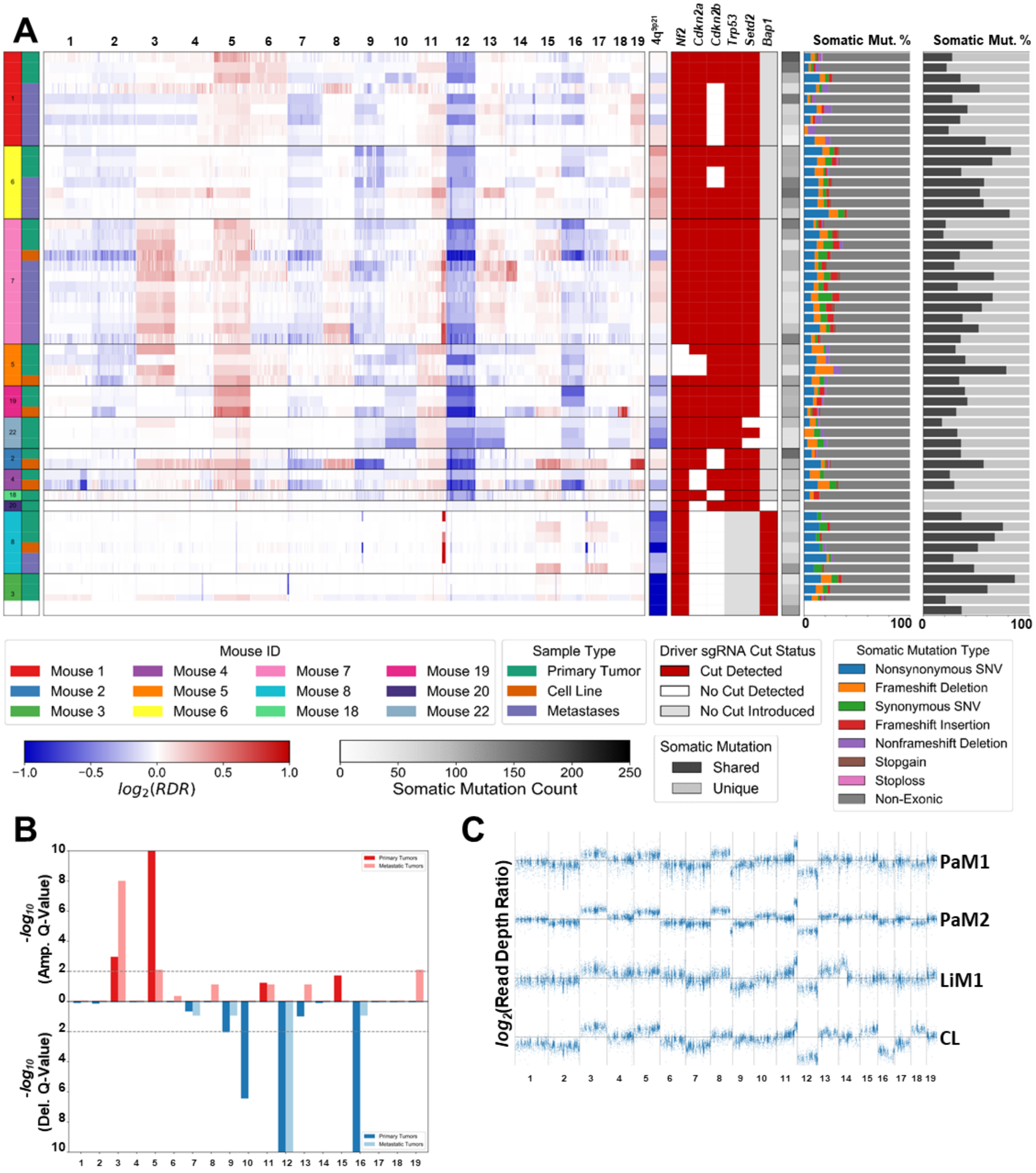


**A)** Summary heatmap displaying WES results. **B)** Clustered bar chart of -log$_{10}$(Q-Value) as determined by GISTIC2 (Methods) displaying the significance of specific focal and broad CNVs in primary and metastatic tumor samples across the experimental cohort. **C)** Representative scatter plots of exon-level log2 (read depth ratios) from MRS (LiM = liver metastasis, PaM = pancreas metastasis, CL = primary tumor–derived cell line).

**Supplementary Fig. S5.**

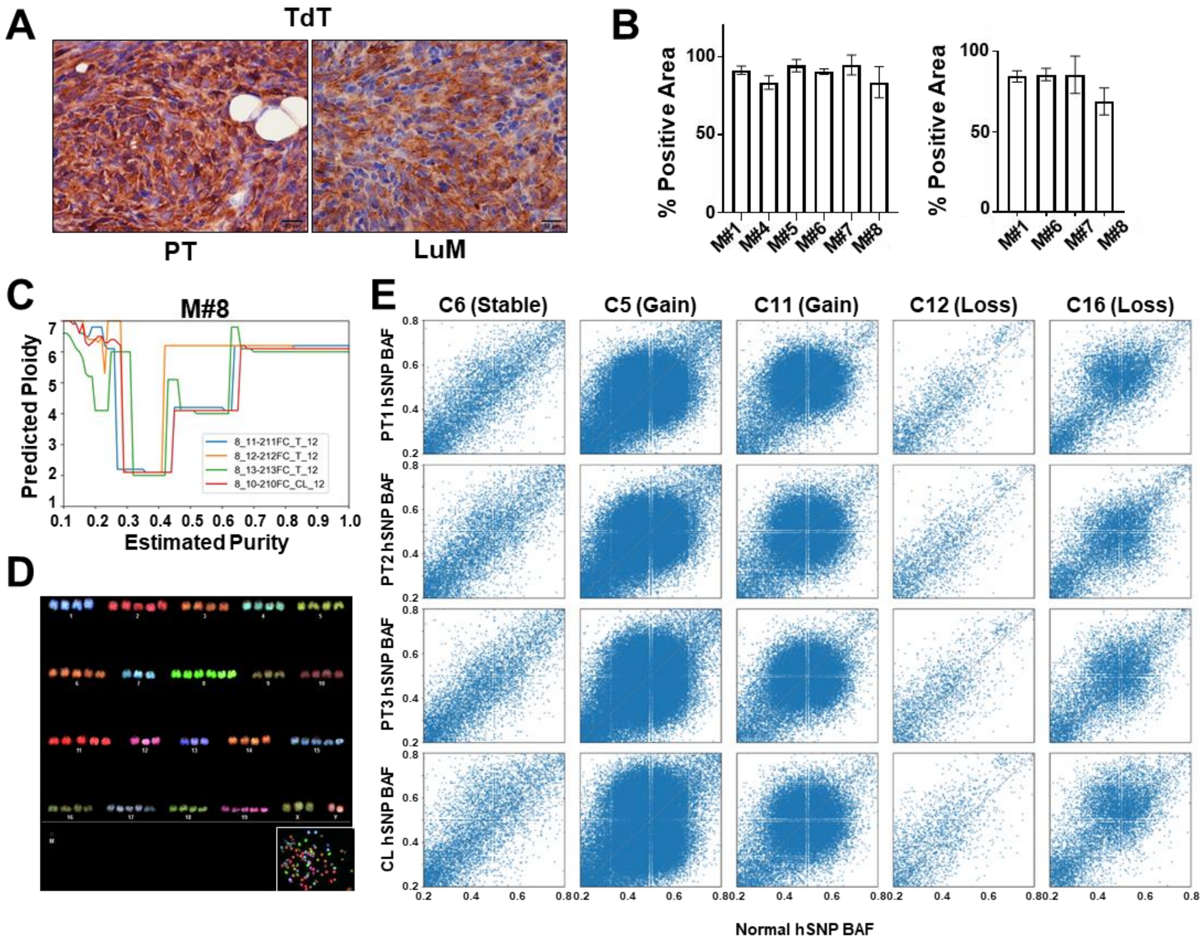


**A)** Representative sections of TdT stained tumor tissues. **B)** Cellularity estimation of primary and metastatic samples as assessed through TdT positive cell quantification (N = 5/group). **C)** Most probable ploidy by log posterior probability at given sample's cellularity as predicted by Sequenza from WGS data (respesentative mouse #8). **D)** Representative spectral karyotyping (SKY) of a short-term culture established from an $Nf2^{KO}$-$Bap1^{KO}$-$4q^{9p21}$ tumor-bearing mouse showing prominent aneuploid features. **E)** Comparison of primary tumor sample and matched normal B-allele frequencies (BAF) of heterozygous SNPs derived from WGS in the matched normal tissue sample ($0.2 \leq$ normal sample SNP BAF $\leq 0.8$). The analysis was performed on chromosomes undergoing gains (5q, 11q) or losses (12q, 16q). A copy-neutral chromosome was used as control (6q). The strong correlation of SNP BAFs between tumor and matched normal samples indicates that these chromosome losses do not cause loss of heterozygosity (LoH) of these SNPs. However, chromosomal loss in diploid genomes would cause LoH of SNPs and the distribution of SNP BAFs in the tumor sample would become bimodal, suggesting that WGD events might precede focal and broad chromosome-level aneuploidy. Error bars represent the standard deviation of technical replicates (B). Scale bar: 100 μm.

**Supplementary Fig. S6.**

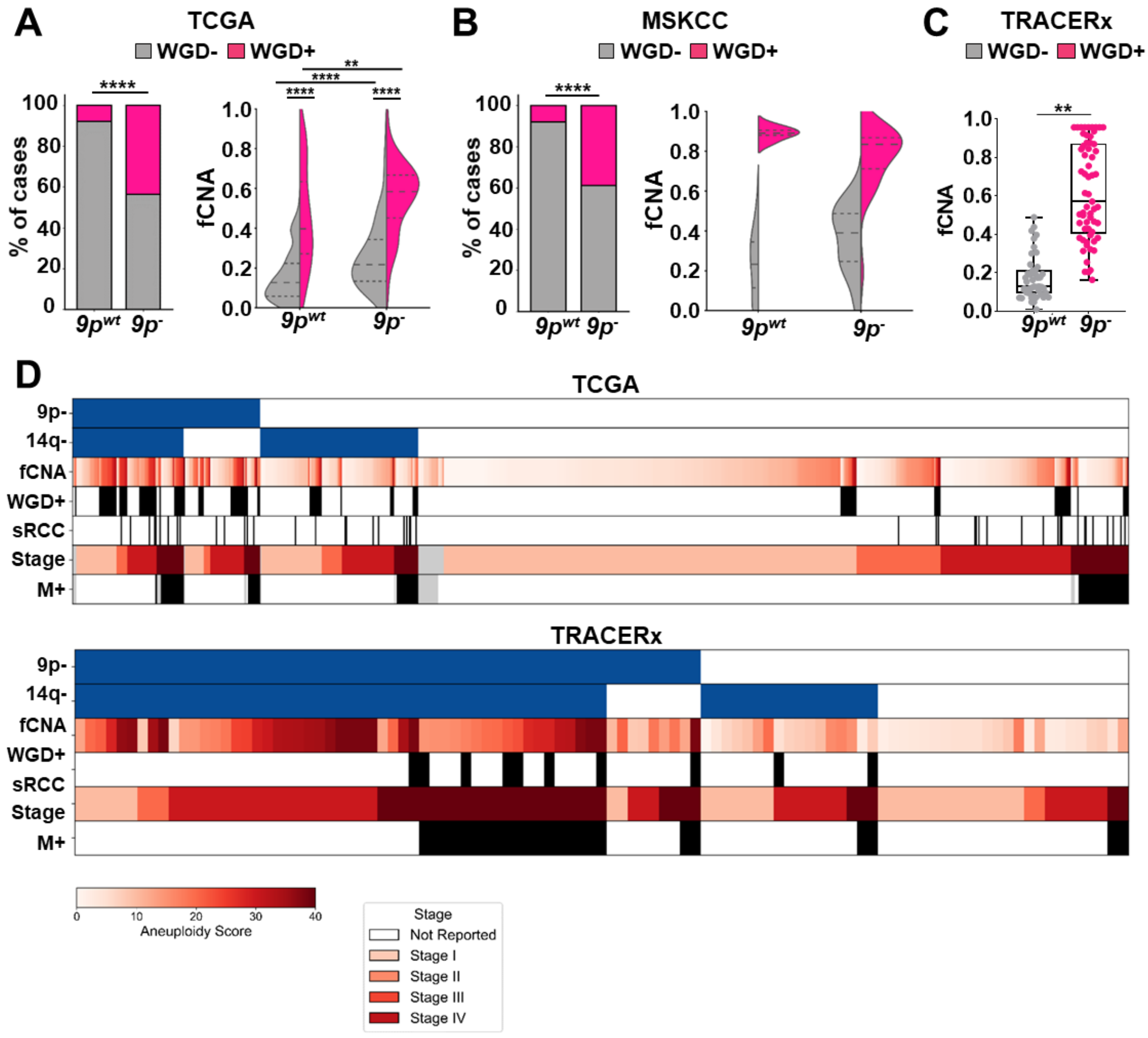


**A)** *Left panel:* Bar chart showing the prevalence of WGD in $9p^{wt}$ and $9p^{-}$ cases in TCGA pan-RCC dataset (N = 788). *Right panel:* Violin plot showing the aneuploidy score distribution as a function of the 9p status and WGD ($9p^{-}$, N = 189; $9p^{wt}$, N = 599). **B)** *Left panel:* Bar chart showing the prevalence of WGD in $9p^{-}$ and $9p^{wt}$ cases in the MSKCC RCC dataset (N = 134). *Right panel:* Violin plot showing the aneuploidy score distribution as a function of the 9p status and WGD ($9p^{wt}$, N = 62; $9p^{-}$, N = 72) in MSKCC RCC cohort. **C)** Box and whisker plot showing the aneuploidy score distribution as a function of the 9p status ($9p^{wt}$, N = 40; $9p^{-}$, N = 61) in TRACERx RCC cohort. **E)** Clinical and genomic annotation of specific features across TCGA pan-RCC cohort (*upper panel*, N = 788) and the TRACERx RCC cohort (*lower panel*, N = 101). Pairwise patients characteristics statistics are in Supplementary table S1. ** $P < 0.01$; **** $P < 0.0001$ by Fisher's exact test (A,B, *left panels*), Mann–Whitney U test (A *right panel*, C). Error bars represent the standard deviation of biological replicates. Dashed lines in violin plot represent median, upper, and lower quartile.

**Supplementary Fig. S7**

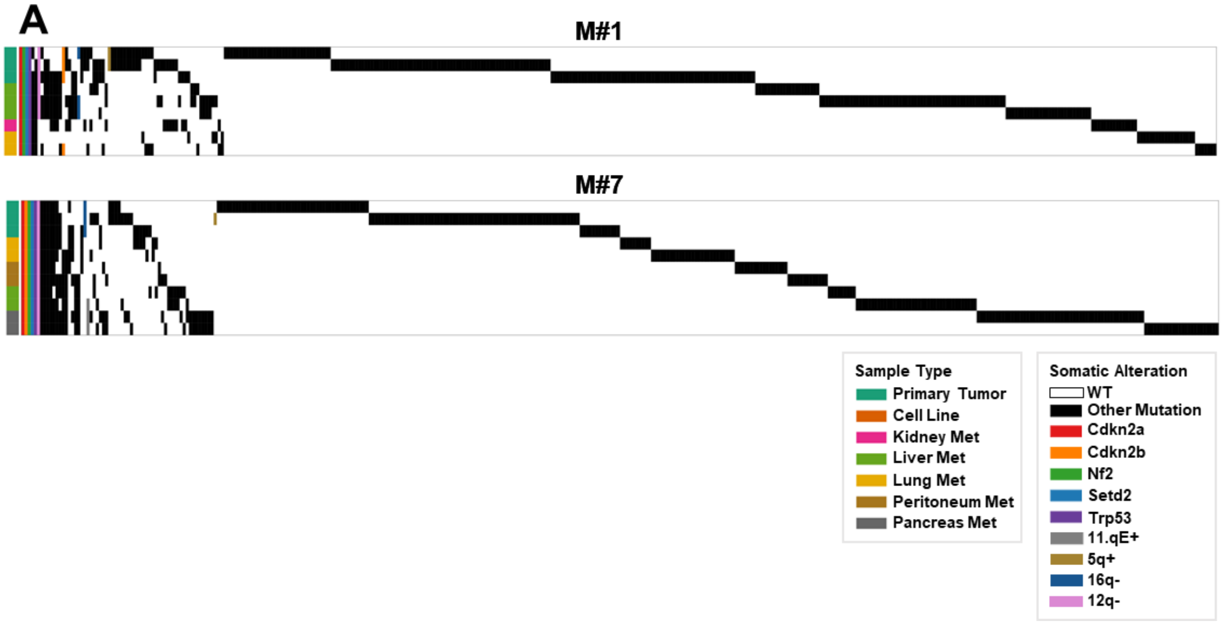


**A)** Patterns of within- and between-lesion heterogeneity among SNVs, CNVs, and indels based on MRS analysis of paired primary tumors and metastases. Early seeding and independent evolution at different sites sampled is readily appreciated.

**Supplementary Table S1.** (see the attached excel file)

Patients characteristics and statistics.
